# Mitoxantrone dihydrochloride, an FDA approved drug, binds with SARS-CoV-2 NSP1 C-terminal

**DOI:** 10.1101/2021.09.14.460211

**Authors:** Prateek Kumar, Taniya Bhardwaj, Rajanish Giri

**Affiliations:** Indian Institute of Technology Mandi, School of Basic Sciences, VPO Kamand, Himachal Pradesh 175005, India

**Keywords:** CD spectroscopy, Fluorescence spectroscopy, Fluorescence anisotropy, Molecular Dynamics Simulations, Drug design

## Abstract

One of the major virulence factors of SARS-CoV-2, NSP1, is a vital drug target due to its role in host immune evasion through multiple pathways. NSP1 protein is associated with inhibiting host mRNA translation by binding to the small subunit of ribosome through its C-terminal region. Previously, we have shown the structural dynamics of NSP1 C-terminal region (NSP1-CTR) in different physiological environments. So, it would be very interesting to investigate the druggable compounds that could bind with NSP1-CTR. Here, in this article, we have performed the different spectroscopic technique-based binding assays of an anticancer drug Mitoxantrone dihydrochloride (MTX) against the NSP1-CTR. We have also performed molecular docking followed by computational simulations with two different forcefields up to one microsecond. Overall, our results have suggested good binding between NSP1-CTR and MTX and may have implications in developing therapeutic strategies targeting NSP1 protein of SARS-CoV-2.

## Introduction

Non-structural proteins (NSPs) of coronaviruses exclusively carry out replication and translation of viral genome, hence they are considered to be one of the prime targets for drug discovery ^1^. SARS-CoV-2 NSP1 is the first protein to be translated as a part of polyprotein 1a and 1ab, and is considered important as it works to suppress the host immune system ^2^. It restricts the host mRNA binding to ribosome by associating with the 40S subunit of host cell using its C-terminal region (CTR) ^3,4^. Upon interaction with ribosome, the NSP1-CTR partially gains α-helical structure propensity ^3,4^. In our previous study, we observed that NSP1-CTR remains in disordered conformation in isolation and is found to adopt a secondary structure in different environments ^5^. It also interacts with mRNA nuclear exporter heterodimer NXF1-NXT1 and hampers the association of mRNA with NXF1, resulting in the abolishment of NXF1 docking on nuclear pore complex ^6^. Recently, a study exposed the detailed insights of host translation inhibition where the eukaryotic translation initiation factors (eIF3 and eIF4F) allosterically control the NSP1 association with ribosome in the early stages of translation in absence of mRNA ^7^. Moreover, mutagenesis-based studies have revealed several key residues in its CTR that are responsible for its interaction with ribosome and abrogation of the translational activity ^4,6^. All these facts make NSP1-CTR an essential drug target that can lead to ultimate blockage of NSP1 activity.

Many studies for drug discovery have been performed in the last few months of pandemic, including the drug repurposing as a time and cost-effective approach ^8,9^. Drug repurposing is the most effective approach to identifying potential drugs against this deadly pandemic-causing disease. The scientific community has tried several existing FDA-approved drugs including different antivirals, antibiotics, steroids, to name a few, against the SARS-CoV-2 infection ^9^. Drugs like remdesivir, tocilizumab, and flavipiravir are being given to the infected patients ^8,10– 13^. In addition, various druggable compounds, including natural compounds, have been reported to be effective against the virus ^14,15^.

One such drug Mitoxantrone dihydrochloride (MTX), has been repurposed and is demonstrated to inhibit the SARS-CoV and SARS-CoV-2 virus entry into cells ^16^. Previously, the antiviral activity of MTX has been established against many other viruses such as Herpes Simplex Virus 1 (HSV-1), and Vaccinia virus ^17,18^. Additionally, the anticancer activity of this drug against lymphoma, breast cancer, and prostate cancer is very well-known ^19–22^. Overall, these properties make it a potential candidate to be tested for its specific binding target with SARS-CoV-2 proteins.

In this report, we have demonstrated the binding of anticancer compound MTX with the C-terminal region of NSP1 (NSP1-CTR) using spectroscopy-based techniques and extensively long computational simulations. According to the finding, MTX binds with NSP1-CTR in its disordered conformation and can be developed into a drug against the virus. Furthermore, this study may provide a useful insight on interacting residues of the C-terminal region for developing antiviral against SARS-CoV-2 NSP1 protein which could prevent its activity inside the host.

## Results

Given the functional importance of NSP1 protein and its disordered C-terminal region, the druggability of NSP1 must be explored to identify novel inhibitors or repurposed drugs against the SARS-CoV-2. Furthermore, the blocking of NSP1-CTR may possibly restrict the binding of NSP1 with ribosomes inside host cells. Following this, we have investigated the binding of an antineoplastic drug MTX with the C-terminal tail of NSP1. Herein, we have performed binding kinetics of MTX with NSP1-CTR using fluorescence and CD spectroscopic techniques. Further, the fluorescence lifetime and anisotropy decay have been measured in the presence and absence of the drug. We have also confirmed the binding through computational modeling and docking studies.

### Fluorescence-based measurements

#### Fluorescence intensity analysis

The binding of MTX with NSP1-CTR is studied experimentally by monitoring the changes in intrinsic fluorescence of NSP1 tryptophan (Trp^161^) residue. As shown in **figure 1A**, the fluorescence intensity of tryptophan residue of the peptide is observed to decrease gradually on increasing the concentration of MTX in the solution. The addition of MTX has substantially quenched the tryptophan fluorescence intensity which indicates the possible binding of MTX with NSP1-CTR. As discussed in previous report on structural dynamics of NSP1-CTR, the _max_ of the tryptophan residue occurs at 346 nm. Therefore, the dissociation constant (K_d_) is obtained by considering the maximum fluorescence intensity at 346 nm using the non-linear fitting of one site-specific binding. The K_d_ for MTX-NSP1-CTR interaction is calculated to be 31.83 µM (R^2^ = 0.9) (**Figure 1B**).

**Figure 1:**
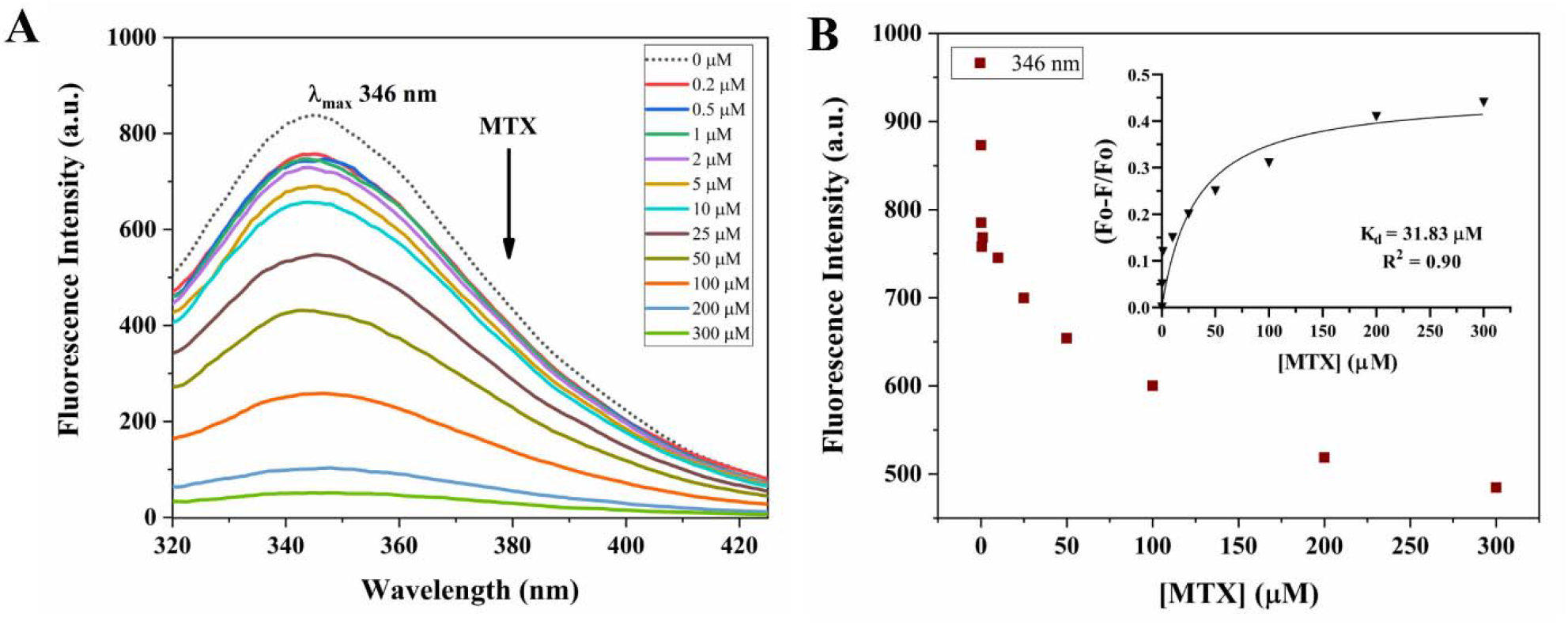
Tryptophan (Trp^161^) fluorescence emission analysis of NSP1-CTR peptide with MTX. **(A)** The decreasing intrinsic fluorescence intensity of NSP1-CTR upon addition of MTX in increasing concentrations (0 to 300 µM). **(B)** The fluorescence intensity at 346 nm is plotted against the concentration of MTX. **Inset:** shows the non-linear fitting of one-site specific binding with the calculated dissociation constant.

#### Fluorescence Lifetime measurements

Generally, the fluorescence decay of tryptophan moiety is susceptible to changes in its surroundings. With conformational changes in proteins due to variation in temperature, viscosity, and surrounding solvent, the fluorescence lifetime changes ^23^. Therefore, the binding and conformational changes in the NSP1-CTR-MTX complex are observed through fluorescence lifetime measurement, where the decay in fluorescence of tryptophan fluorophore was recorded. As observed in graphs depicted in **figures 2A** and **2B**, the average fluorescence lifetime of Trp^161^ of NSP1-CTR shows a decreasing trend compared to the fluorescence lifetime in unbound conformation. The average lifetime of unbound protein is measured to be 1.98 ns with a chi-square (χ^2^) value of 1.11, whereas, in presence of 200 µM MTX, the average lifetime decreased to 1.66 ns (χ^2^ = 1.08) (see **table 1**). The declining fluorescence lifetime is evident in the binding of MTX with NSP1-CTR peptide.

**Figure 2:**
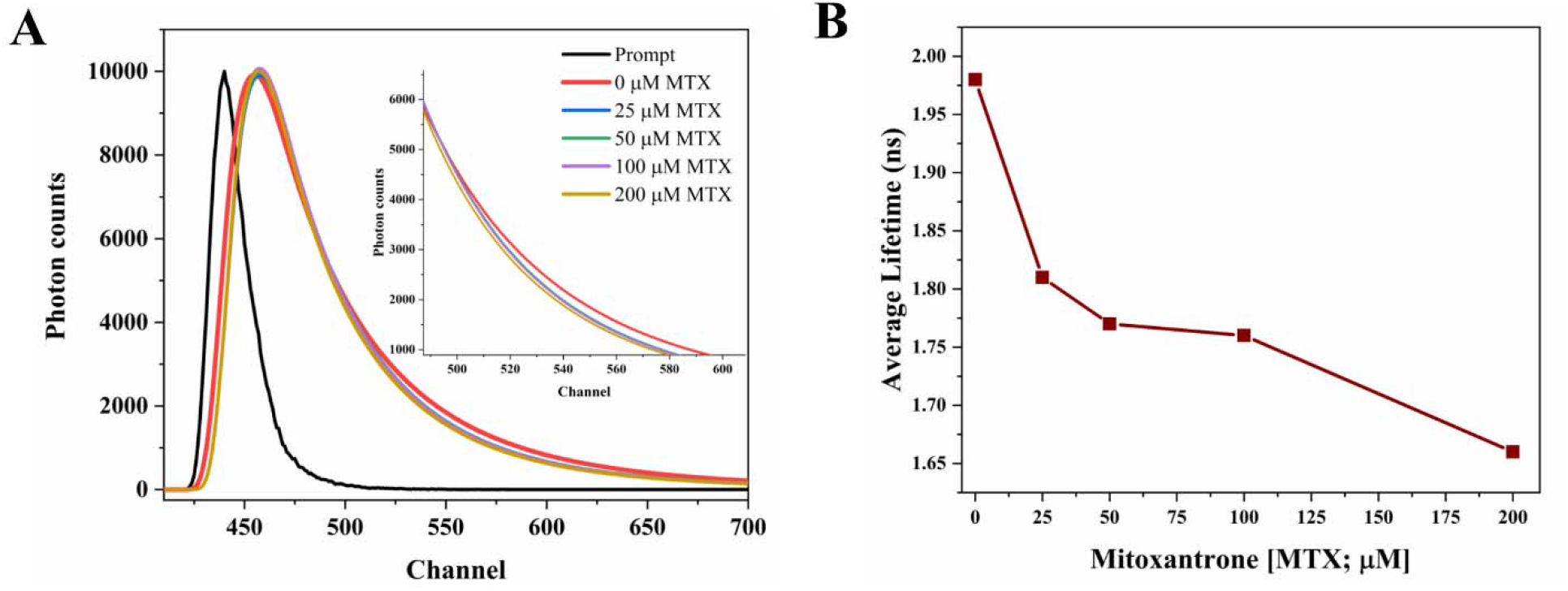
Assessment of Trp^161^ by using fluorescence lifetime measurement. **(A)** Fluorescence lifetime decay curve of NSP1-CTR peptide in absence and presence of drug MTX at four increasing concentrations. **(B)** Illustration of decrease in average fluorescence lifetime of Trp^161^ with increasing concentration of MTX.

**Table 1:** Decreasing average fluorescence lifetime of Trp^161^ of NSP1-CTR upon addition of MTX.

| Mitoxantrone Concentration (μM) | $\tau_1 \pm s$ (ns) | $\tau_2 \pm s$ (ns) | $\tau_3 \pm s$ (ns) | $\tau$ (average lifetime, ns) | $\chi^2$ - value |
| --- | --- | --- | --- | --- | --- |
| 0<br>(Only NSP1-CTR; 7.5 μM) | $2.42 \pm 0.10$ | $0.61 \pm 0.02$ | $5.28 \pm 0.04$ | 1.98 | 1.11 |
| 25 | $2.31 \pm 0.06$ | $5.12 \pm 0.06$ | $0.63 \pm 0.02$ | 1.81 | 1.06 |
| 50 | $2.23 \pm 0.08$ | $5.02 \pm 0.05$ | $0.59 \pm 0.02$ | 1.77 | 1.13 |
| 100 | $2.26 \pm 0.08$ | $4.96 \pm 0.06$ | $0.62 \pm 0.02$ | 1.76 | 1.24 |
| **200** | $2.24 \pm 0.08$ | $4.91 \pm 0.06$ | $0.58 \pm 0.01$ | 1.66 | 1.08 |

\*The quoted errors are sample standard deviations as $\pm s$ .

#### Time-resolved Anisotropy decay analysis

As aforementioned, tryptophan fluorescence is susceptible to changes in its local environment. Its global and local rotations are affected by the changes in the surroundings. Therefore, to further observe the binding of MTX with NSP1-CTR, we have performed an anisotropy decay analysis of Trp^161^ residue. Here, the fluorescence decays of tryptophan residues in perpendicular and parallel emission polarization of light are recorded. The decay curve is fitted using a bi-exponential decay function which defines the fast (**θ**_1_) and slow (**θ**_2_) rotational correlational time where **θ**_1_ is the local rotation of Trp^161^ while the **θ**_2_ is the global rotation of the peptide ^24^. In unbound state, NSP1-CTR (30 µM) has an average rotational correlation time (τ_r_) equals to 4.46 ns (χ^2^ = 1.19) **(Figure 3A & 3B)**. Interestingly, after binding with MTX (25 µM), its average rotational correlation time decreases to 962.1 ns (χ^2^ = 1.14) **(Figure 3C & 3D)**. (**θ**_1_) and (**θ**_2_) for NSP1-CTR have been calculated to be 1.13 ns and 15.6 ns respectively, while in presence of MTX, the (**θ**_1_) and (**θ**_1_) are observed to be 1070.1 ns and 1.13 ns. These results correspond to the interaction of the drug with the peptide as binding of MTX has slowed the rotation of the tryptophan residue and the peptide.

**Figure 3:**
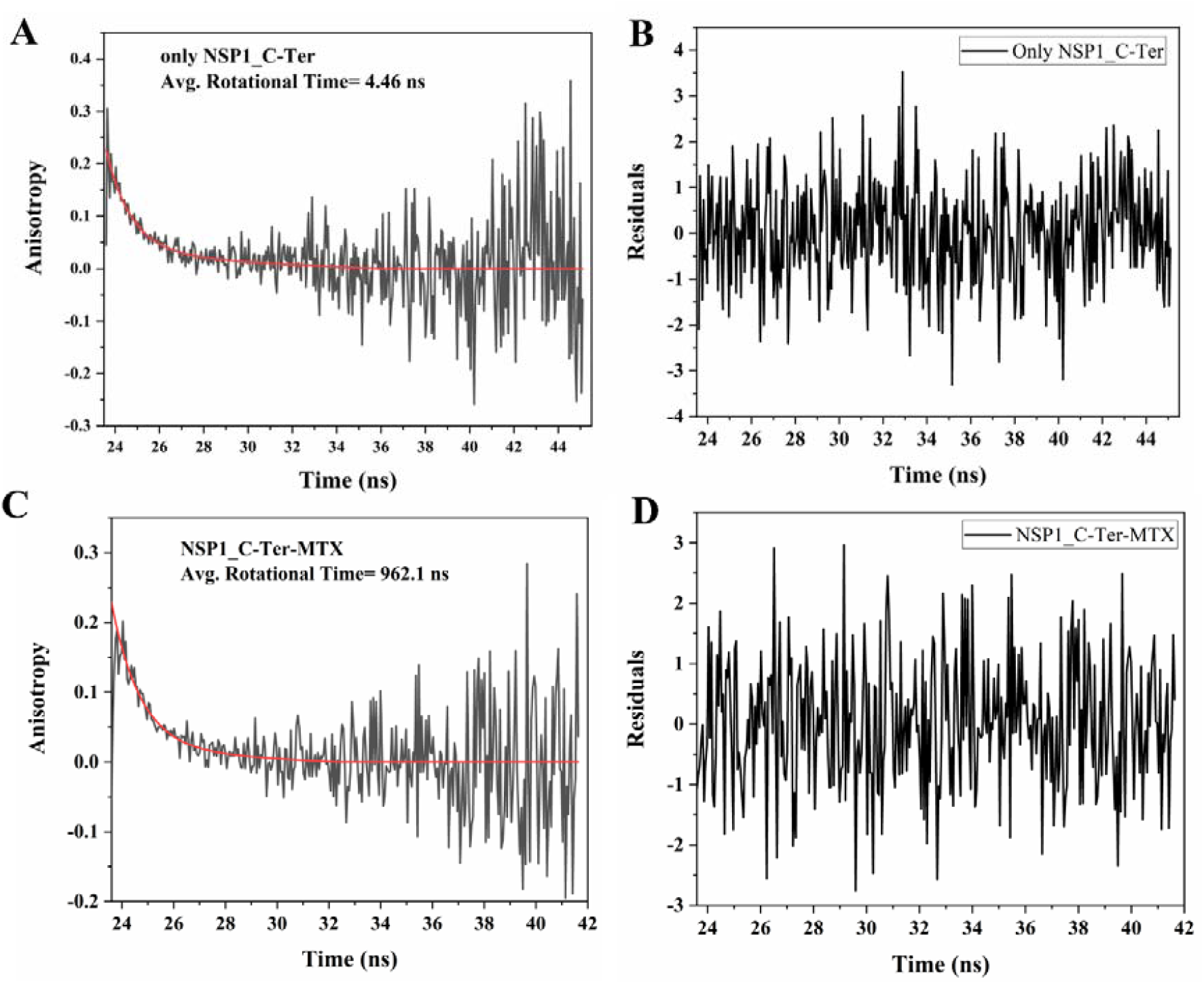
Fluorescence anisotropy decay measurements of NSP1-CTR. (**A**) the conformational dynamics in unbound state of NSP1-CTR (30 µM), and (**B**) in the bound state with compound MTX (25 µM). Figure (**C**) and (**D**) depict the residual of the fitting curve corresponding to unbound and bound state of NSP1-CTR.

### Circular dichroism-based structural alterations in NSP1-CTR

In order to gain insights into changes in the secondary structure of NSP1-CTR by virtue of MTX interaction, we employed CD spectroscopy and investigated the effect of the binding of MTX on its structure. The far-UV CD spectra of NSP1-CTR were recorded is presence and absence of MTX. As reported before, NSP1-CTR remains in a disordered state in isolation ^5^. Therefore, in the unbound state, the negative ellipticity at ∼200 nm wavelength represents a typical spectrum corresponding to the disordered state of NSP1-CTR. However, the presence of MTX has only caused minimal fluctuations in the negative ellipticity at ∼200 nm (**Figure 4**). However, these CD results demonstrate that in presence of the drug MTX does not cause any noticeable structural changes in the secondary structure of the peptide, but MTX may bind with NSP1-CTR in the latter’s disordered form.

**Figure 4:**
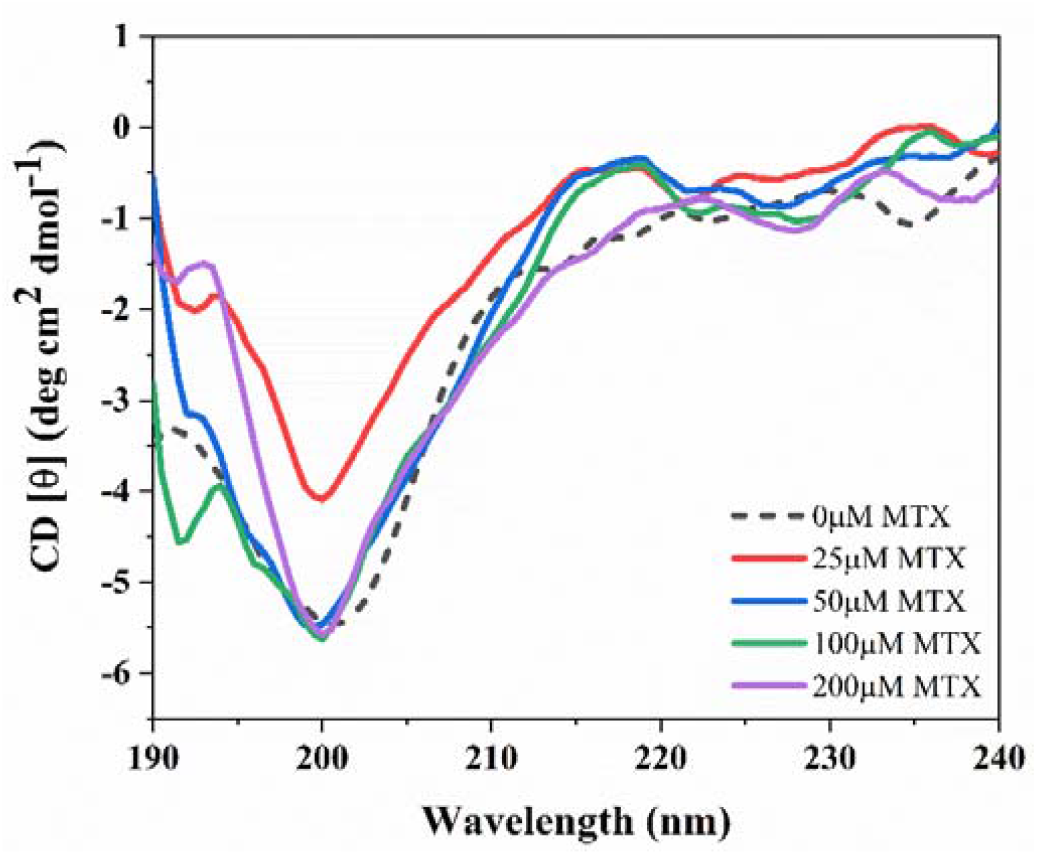
Circular Dichroism spectra of NSP1 C-terminal upon interaction with compound MTX on varying concentrations from 0 to 200 µM in comparison with the unbound state of NSP1-CTR at 25 µM.

### Molecular docking analysis

As previously reported by our group, the NSP1-CTR is mainly in disordered conformation and contains helical structure forming tendencies in different environmental conditions ^5^. However, it gains alpha helix upon interacting with its binding partner i.e., 40S ribosomes (PDB ID: 6ZOJ). MD simulations of its modeled structure have also revealed the α-helical structure, therefore, we have selected the structure at last frame (at 500 ns; described as NSP1-F1 in our previous study) of NSP1-CTR from the simulation trajectory in aqueous conditions ^5^.

The MTX drug is docked against NSP1-CTR by enclosing the whole structure as a receptor grid, as the structure constitutes no proper binding pocket. *Glide* scoring function of the Schrodinger suite is employed for docking and Prime module for MM/GBSA calculation. The MTX has scored a docking score of -5.335 kcal/mol and binding energy of -47.49 kcal/mol. Besides, the strong binding with six hydrogen bonds and two salt bridge interactions has been observed between NSP1-CTR and MTX (**Figure 5**). According to binding pose and ligand interaction diagram, the residues involved are Ser12 (or Ser142; according to full-length structure), Asp14 (or Asp144), Arg41 (or Arg171), and Glu25 (or Glu155).

**Figure 5:**
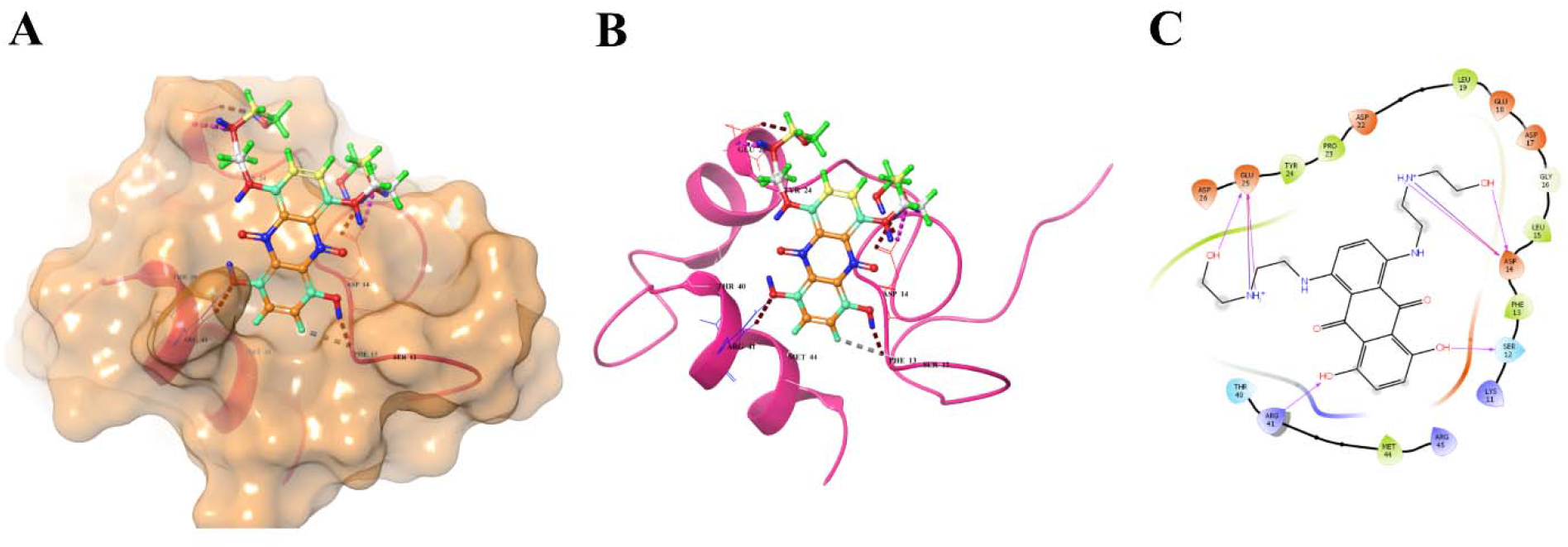
In-silico interaction analysis: Molecular docking of Mitoxantrone against NSP1-CTR by enclosing whole structure in the grid. **(A)** The surface representation of NSP1-CTR with docked drug MTX, **(B)** 3D cartoon representation of protein-ligand binding pose, and **(C)** 2D interaction showing interacting residues of NSP1-CTR with MTX is depicted.

### MD Simulations analysis

Aiming to validate the experimental results showing the binding of MTX with NSP1-CTR peptide, we further conducted the simulation of the docked complex using two different forcefields and examined the interaction stability through computational means.

Firstly, the MD simulations are performed upto 1 µs with OPLS 2005 forcefield, which is implanted in Desmond package and Schrodinger’s Maestro interface. According to our microsecond timescale data, the majorly disordered peptide of NSP1-CTR has shown significantly good interactions during the entire simulation period except for few frames. Based on the RMSD, the MTX bound NSP1-CTR has shown an average RMSD of ∼7Å in 1 µs trajectory wherein initial 300 ns time, some fluctuations in RMSD are observed which were then stabilized till end of simulation time **(Figure 6A; upper panel)**. Consequently, the compactness parameter (Rg) of NSP1-CTR has also shown high variation in the simulation period, possibly due to its disordered nature **(Figure 6A; middle panel)**. Also, the number of hydrogen bonds has been affected due to some fluctuations and averaged out to approx. 2 for 1 µs long simulation trajectory **(Figure 6A; lower panel)**. As shown in **figure 6B**, the fluctuations in residues have been observed till 5Å for middle regions while higher for terminal regions. The reason could be its intrinsic property to bind with compound MTX in its disordered state which is above shown through CD spectroscopic analysis.

**Figure 6:**
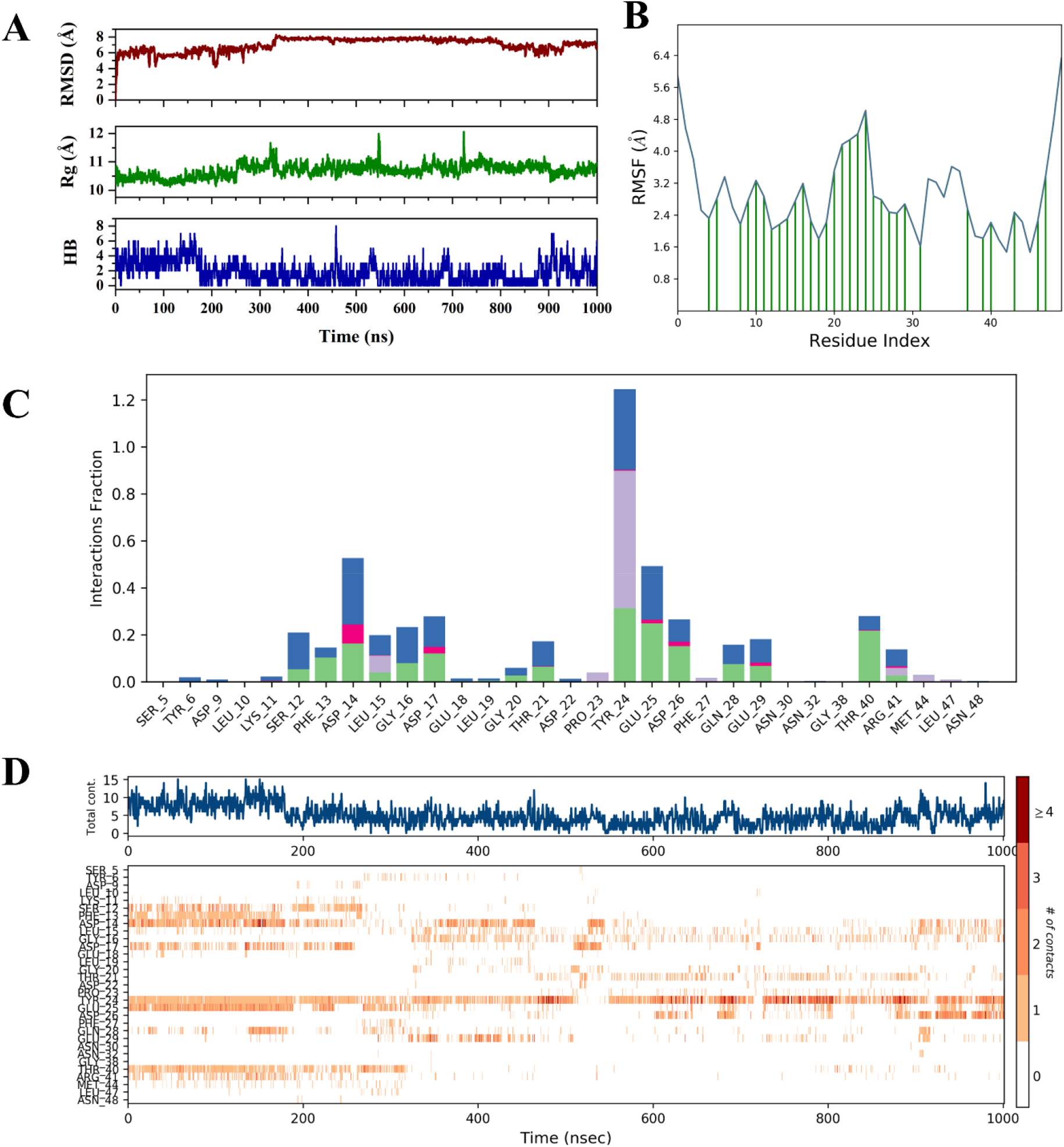
Molecular dynamics simulation analysis of MTX bound NSP1-CTR using OPLS 2005 forcefield: **(A)** RMSD, Rg, of C-alpha (C_α_) atoms and hydrogen bonds (from up to down), (**B)** RMSF analysis highlighting interactions of NSP1-CTR formed with MTX, (C) Histogram plot for representation of the fraction of interaction with residues throughout the simulations where hydrogen bonds (green), water bridges (blue), hydrophobic (purple), and ionic (red) are shown, and one bar with multiple colors shows multiple interactions with that specific residue and (D) Timeline representation of interacting residues where dark color depicts more number of interactions.

Further, this is also evident from the timeline representation of secondary structure analysis of 1 µs simulation trajectory where the majority of secondary structure element is disordered in nature (**Supplementary figure 1**). In last, the contacts made by MTX with residues of NSP1-CTR have been observed in a significant proportion. A few residues like Tyr24 (or Tyr154) and Asp14 (or Asp144) interact with multiple bonds for most of the simulation time **(Figure 6C & 6D)**. The structural dynamics and interaction of MTX with NSP1-CTR are shown in **supplementary movie 1**.

Another forcefield for examining the protein-ligand complex stability is chosen as GROMOS54A7 in Gromacs simulation package. The outcomes are in good correlation with the above simulation of longer time period. As shown in **figure 7A (upper panel)**, the RMSD of the complex remains stable throughout the simulation period, with only a few noticeable fluctuations in RMSD values that are observed after 50 ns to 250 ns. The simulation trajectory has attained stability after 250 ns with an average RMSD value of 0.75 nm. In view of compactness, the MTX bound NSP1-CTR has attained an average Rg value of 1.05 nm (**Figure 7A; middle panel)**. Accordingly, the fluctuation in residues was also observed in terms of RMSF that is found to be varying between 0.4-0.6 nm during most of the regions. However, terminal regions have shown high flexibility as they do not constitute any structure (**Figure 7A; lower panel)**. The average number of hydrogen bonds are calculated to be approximately 2 for the simulation trajectory (**Figure 7B)**. Also, the PCA calculation is done for MTX bound NSP1-CTR, where it has shown a little scattered cluster with the distance of -2 to 2 nm (**Figure 7C & 7D)**. The change in secondary structure is also observed concerning time where no significant change is observed, and the peptide remains majorly disordered **(Supplementary figure 2)**. Overall, both simulation results are in agreement with the experimental binding of MTX with NSP1-CTR within a favorable range.

**Figure 7:**
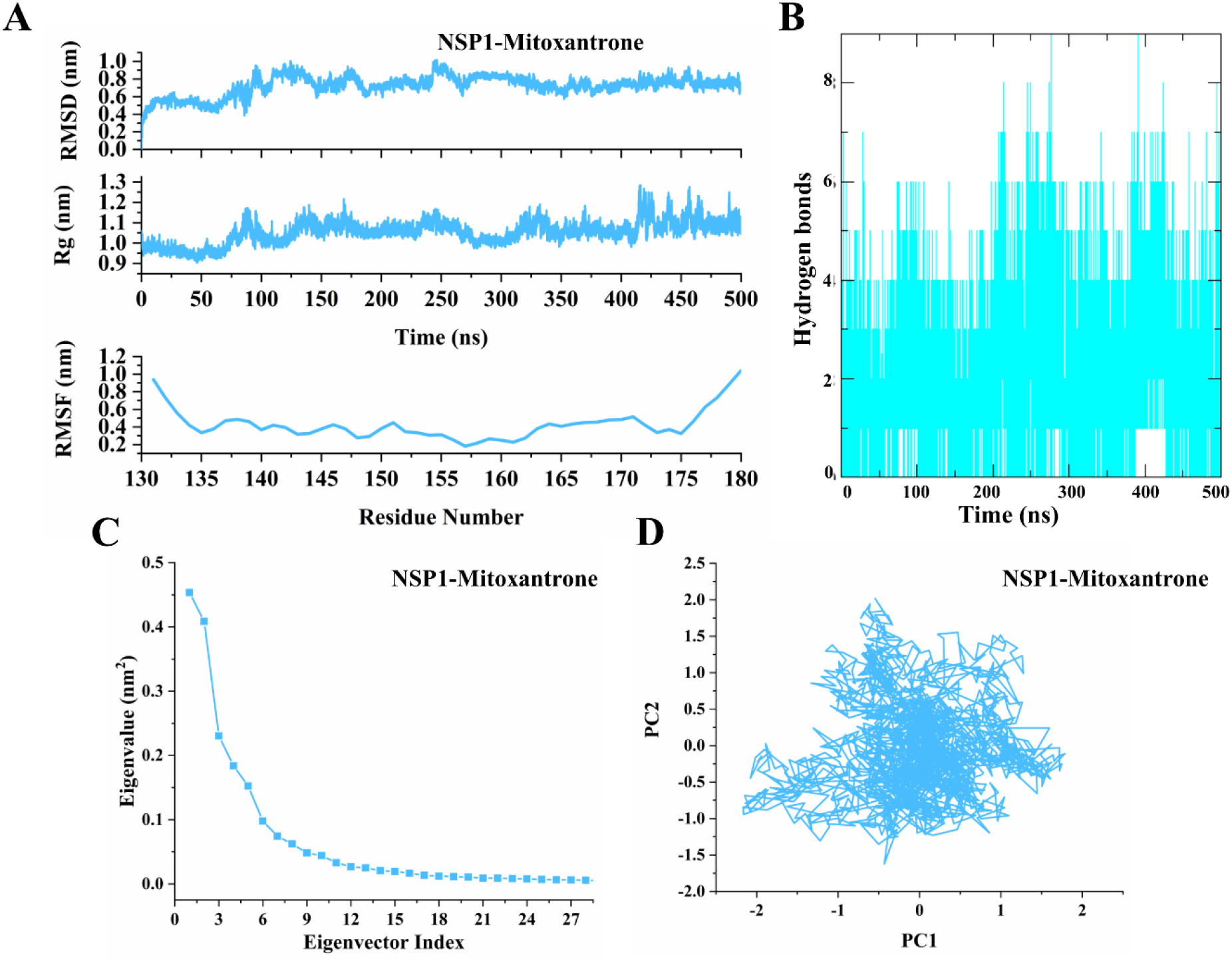
Molecular dynamics simulation analysis of MTX bound NSP1-CTR using GROMOS 54A7 forcefield: **(A)** RMSD, Rg, and RMSF (from up to down), (**B)** hydrogen bonds analysis, **(C)** Eigenvector vs. Eigenvalue plot, and **(D)** principal component analysis of last 20 ns simulation trajectory.

## Discussion and Conclusion

Since the COVID-19 pandemic began, a plethora of studies on the repurposing of existing drugs and the identification of new drugs have been reported. However, none of the drugs were successful after clinical trials with high efficiency and fewer side effects. Several antivirals like Remdesivir, Flavipiravir, Lopinavir, etc., antimalarial drug Hydroxychloroquine have been tested in clinical trials as well as these have been used earlier with emergency usage authorization. These drugs have been studied against multiple target proteins of SARS-CoV and SARS-CoV-2. Among various important druggable targets of SARS-CoV-2, NSP1 protein is largely responsible for host immune response suppression. Therefore, we have studied the structure-based drug discovery against its multifunctional domain, C-terminal region, through spectroscopic and computational approaches. Here, we have investigated the binding kinetics of an anticancer compound, MTX, against NSP1-CTR.

Initially, MTX was developed as an analog of doxorubicin, an anthracycline, and is currently being used as an anticancer drug since the 1980s due to its better antitumor activity and less cardiotoxic properties ^20^. The antiviral activity of this drug has also been established in several viruses. Recently, Zhang et al. has observed the role of heparan sulfate, which is present on the cell surface, as a binding mediator of ACE-2 assisted attachment of SARS-CoV and SARS-CoV-2 in host cells. Further, they have abolished that interaction using the drug MTX ^16^. It targets the heparan sulfate on host cells which ultimately prevents the viral entry ^16^. In Vaccinia Virus, it prevents viral maturation and assembly by blocking the processing of mature structural proteins ^18^. Another study on Herpes simplex virus 1 (HSV-1) reported the binding of MTX has suppressed the viral gene expression ^17^. Moreover, a report based on computational docking has shown that MTX has a higher docking score against SARS-CoV-2 main protease among various other drugs ^25^.

NSP1-CTR is a crucial factor that suppresses the host immune system with its disorder to order transition properties inside the ribosome subunit, which ultimately restricts the mRNA from performing the translation. In this manuscript, we have performed experimentations on NSP1-CTR with literature identified an anticancer drug, Mitoxantrone. The synthetic peptide of NSP1-CTR has a single Trp at 161^st^ position, monitored for fluorescence quenching and lifetime measurements in presence of the chosen compound MTX. The compound MTX has exerted a docking score of more than -5.0 kcal/mol and significantly good binding energy (MM-GBSA score). Its interaction with key residues like Ser142, Asp144, Lys164, Arg171 is also observed. Taking these results together, it is evident that MTX binds NSP1-CTR with excellent efficiency. To further check the structural changes in NSP1-CTR upon binding with MTX, we performed CD spectroscopy at different concentrations of MTX. The observation suggests that MTX does not change any conformation of NSP1-CTR. Based on simulations with two different forcefields, it is also clearly evident that the NSP1-CTR interacts with compound MTX in its disordered state as no structure gain property is observed in virtual simulations similar to experimental observation through CD spectroscopy. Due to the unavailability of full-length NSP1 or the N-terminal region, the binding experiments couldn’t be performed, which may or may not have any implications based on the current study.

Besides, we have also performed docking calculations on 665 compounds from the PubChem database, similar to Mitoxantrone or having anthracene as their parent molecule. As per our observations, a total of 33 compounds have been identified with docking scores more than Mitoxantrone (−5.3 kcal/mol). The compound with Pubchem ID: 88654295 ((2R)-2-Amino-N-[2-[[4-[2-[[(2R)-2-amino-3-sulfanylpropanoyl]amino]ethylamino]-5,8-dihydroxy-9,10-dioxoanthracen-1-yl]amino]ethyl]-3-sulfanylpropanamide), has highest docking score of -6.9 kcal/mol amongst all other compounds and have multiple interactions with Ser12 (or Ser142), Asp14 (Asp144), Asp17 (Asp147), Glu18 (Glu148), Glu25 (Glu155), Gln28 (Gln158), Thr40 (Thr170), and Arg41 (Thr171) (**Supplementary figure 3**). The resultant compounds with docking scores and their structures are shown in the supplementary file (**Supplementary figure 4A & 4B and supplementary table 1**). Furthermore, we have also checked the interaction of around 160 similar conformers of Mitoxantrone from the PubChem database. All these conformers are anthracene molecules with the same backbone and different functional groups. In molecular docking of these compounds, most have shown similar interactions and docking score of maximum -6.1 kcal/mol. In **supplementary table 2**, we have listed the top 15 identified compounds with docking score >-5.3 kcal/mol and their 2D structures shown in **supplementary figure 5**.

Based on these observations, it is highly plausible that the compounds like MTX and others with anthracene moiety could be explored further for experimental investigation against different targets of SARS-CoV-2. Additionally, the interactions of MTX and other similar compounds with key residues of NSP1-CTR also confer that it may inhibit the functions of NSP1 and restrict its association with the ribosomal subunit. Therefore, in the future, based on this study, it is highly possible to target NSP1 and, specifically, its CTR for developing therapeutic strategies, and further experimentation can be done using MTX as a lead or parent molecule.

## Material and Methods

### Materials

#### NSP1 peptide

The sequence of NSP1-CTR (residues 131-180) “NH2-AGGHSYGADLKSFDLGDELGTDPYEDFQENWNTKHSSGVTRELMRELNGG-COOH” was retrieved from the UniProt (ID: P0DTC1.1). The peptide of purity >81 % was ordered from GenScript USA.

#### Compound

Mitoxantrone (1,4-dihydroxy-5,8-bis[2-(2-hydroxyethylamino) ethylamino] anthracene-9,10-dione; dihydrochloride) was purchased from Sigma-Aldrich, USA.

### Fluorescence measurement

Intrinsic tryptophan fluorescence of NSP1-CTR (Trp^161^) was monitored using a spectrofluorometer (Horiba Scientific Fluorolog, model no. 1073) with an excitation wavelength of 284 nm from 300 nm to 450 nm wavelength range at 25°C. The slits bandwidths for both excitation and emission were kept at 4 nm. Binding of MTX with NSP1-CTR was performed by titration experiment where protein concentration was kept constant at 8 μM in presence of 50 mM sodium phosphate (pH 7.4) buffer. The inner filter effect in fluorescence data was corrected using the following equation ^23^:

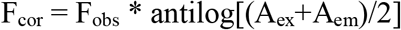

where F_obs_ and F_cor_ are measured and corrected fluorescence intensities, respectively. A_ex_ and A_em_ represents the absorbance of the sample at excitation and emission wavelengths, respectively.

### Fluorescence lifetime measurement

The fluorescence lifetime of tryptophan residue (Trp^161^) in NSP1-CTR in absence and presence of MTX was measured using the Horiba Scientific DeltaFlex TCSPC system at 25°C. The samples were excited at 284 nm using the nanoLED as a light source. The wavelength of the emission monochromator was set at 346 nm. The measurement range was adjusted to 400 ns with bandpass of 32 nm and peak preset of 10000 counts. Ludox was used to correct the instrument response factor (IRF) at 284 nm. The concentration of NSP1-CTR was kept constant at 8 μM and in presence of 50 mM sodium phosphate (pH 7.4) buffer. Finally, the three exponential decay function fitting was used to obtain the tryptophan lifetimes.

### Circular dichroism (CD) measurement

Far-UV CD spectra of NSP1-CTR in 50 mM sodium phosphate (pH 7.4) buffer were recorded using Jasco-1500 spectrophotometer from 190 nm – 240 nm, using 1 mm quartz cuvette (Hellma, Plainview, NY, USA) at 25°C. The bandwidth was kept at 4 nm for all measurements. For CD spectra of NSP1 C-terminal peptide, a control spectrum consisting of only buffer was subtracted from the peptide spectrum. To analyze the CD spectra of peptide in presence of compounds, CD spectra of compounds were subtracted from the peptide spectrum. For all measurements, the protein concentration was kept at 25 μM. All the measurements were smoothened by 5 points through the adjacent-averaging method in Origin software.

### Time-resolved Anisotropy decay measurement

To confirm the binding of NSP1-CTR and MTX, time-resolved anisotropy decay was measured using DeltaFlex TCSPC system (Horiba Scientific) at 25°C. A 1 ml pathlength quartz cuvette was used to record the measurements where the measurement range was set up to 200 ns. The bandpass of 16 nm, peak preset of 1,000 counts, and repetition rate of 1 MHz were used during experimentation. The protein concentration was kept constant at 30 μM. At 284 nm wavelength, ludox was used for adjusting the instrument response factor (IRF). Using the data analysis software DAS6 (Horiba Scientific), the resultant anisotropy decay curves were fitted into a bi-exponential decay function ^23^:

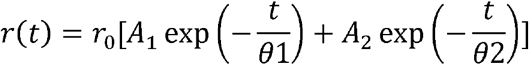

where, anisotropy at time *t* is denoted by *r*(*t*) and initial anisotropy is denoted by *r*_0_, and *A*_1_ and *A*_2_ defines the rotational correlation time □_1_and □_2_, respectively.

### Molecular Docking

The docking of MTX against the simulated structure of NSP1-CTR was performed and the molecular interactions between the complex were analyzed using the Glide XP (Extra Precision) program in the Schrodinger suite ^26^. The parameters for protein structure and ligand preparation are kept as previously reported ^27^. Grid coordinates were set as 28.0, 24.0, and 33.18 Å for X, Y, and Z, respectively. The binding energy of complex was calculated using Molecular Mechanics-Generalized Born Surface Area (MM-GBSA) approach embedded in Prime module of Schrodinger ^28^.

### Molecular Dynamics Simulations

The MD simulations were performed to observe the binding of MTX with NSP1-CTR using two different forcefields viz. OPLS 2005 forcefield in Desmond simulation package (v2018.4) ^29^ and GROMOS54A7 forcefield, utilizing the in-house cluster facility, simulation of the complex was achieved in Gromacs v5.1.2 ^30^.

We have used our previously reported protocol for simulations of protein-ligand complex using Desmond ^31^. Briefly, we have utilized TIP3P water model along with counterions present in the system for charge neutralization. Nose-Hoover and Martina-Thomas-Klein methods were used for temperature and pressure controlling in equilibration and final production runs. A total of one microsecond (µs) simulation was performed. For MD simulations in Gromacs, the SPC water model was added and charge neutralization was done by adding counterions in the simulation setup. The energy minimization was performed by implementing the 50,000 steps of steepest-descent algorithm with Verlet cut-off scheme to calculate the neighboring interactions. Further, equilibration process was carried out under NPT and NVT conditions for 1ns. Parrinello-Rahman and V-rescale methods were utilized for pressure and temperature coupling, respectively. The

Gromacs topology of the compound was generated using PRODRG server ^32^. LINCS algorithm was used for calculating bond parameters ^33^. Furthermore, the simulation analysis parameters are analyzed using gmx toolbox, including rms, rmsf, gyrate, hbond, covar, and anaeig commands. For visualization, UCSF Chimera ^34^ and Schrodinger Maestro were used.

## Authors’ contribution

RG: Conception, design, and review of the manuscript. PK, and TB: acquisition and interpretation of data, writing of the manuscript. PK and TB have contributed equally.

## Acknowledgments

All the authors would like to thank IIT Mandi and HPC utility for providing facilities. RG would like to acknowledge the Department of Biotechnology, Govt of India (BT/11/IYBA/2018/06) and SERB-India (CRG/2019/005603). TB is grateful to the Department of Science and Technology, Govt. of India, for her INSPIRE fellowship for Funding.

