## supplementary information for "Mitoxantrone dihydrochloride, an FDA approved drug, binds with SARS-CoV-2 NSP1 C-terminal"

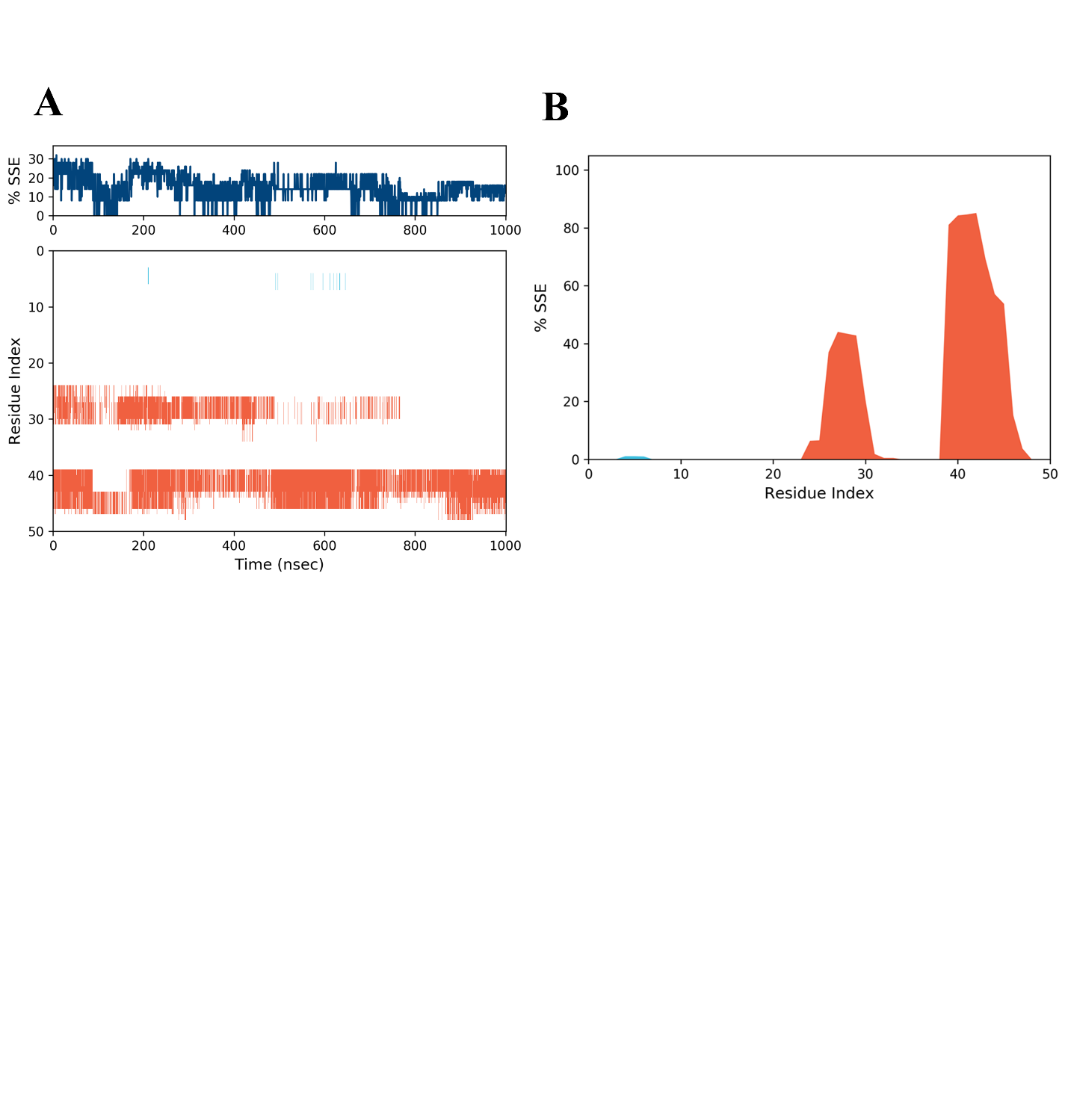

**Supplementary Figure 1:** Illustration of secondary structure change in NSP1-CTR in presence of compound, Mitoxantrone. (A) Timeline representation of each residue of forming helical and beta sheets in respective frame of one microsecond long simulation trajectory. (B) Total secondary structure element (%SSE) is shown for each residue during entire simulation period. The orange color shows helical region and cyan shows the beta sheets.

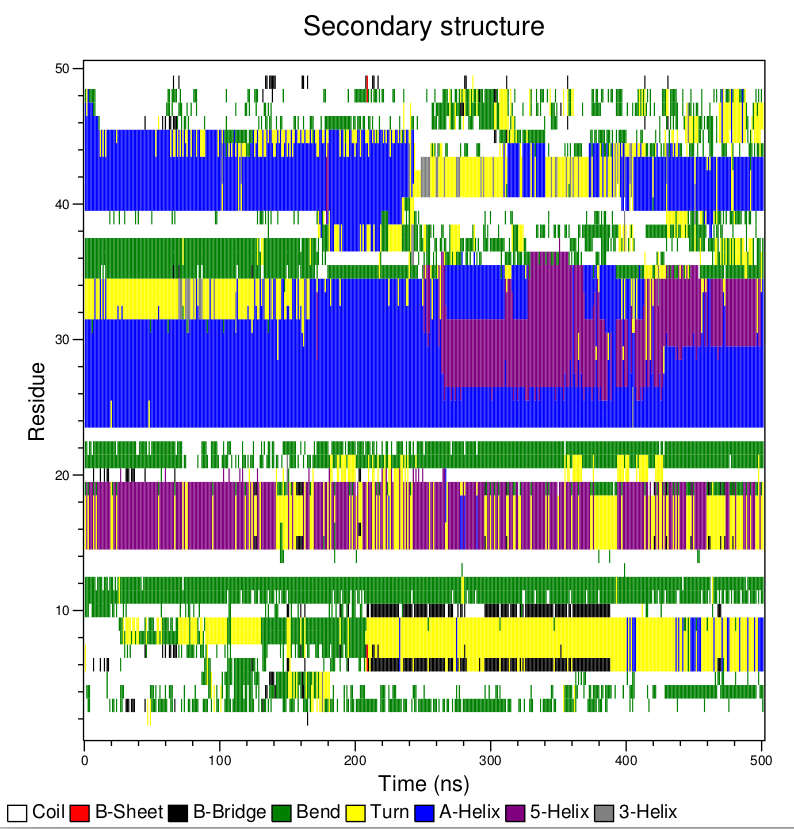

**Supplementary Figure 2:** Timeline representation of secondary structure change in NSP1-CTR during 500 ns long simulation trajectory. The colors are illustrated within the figure.

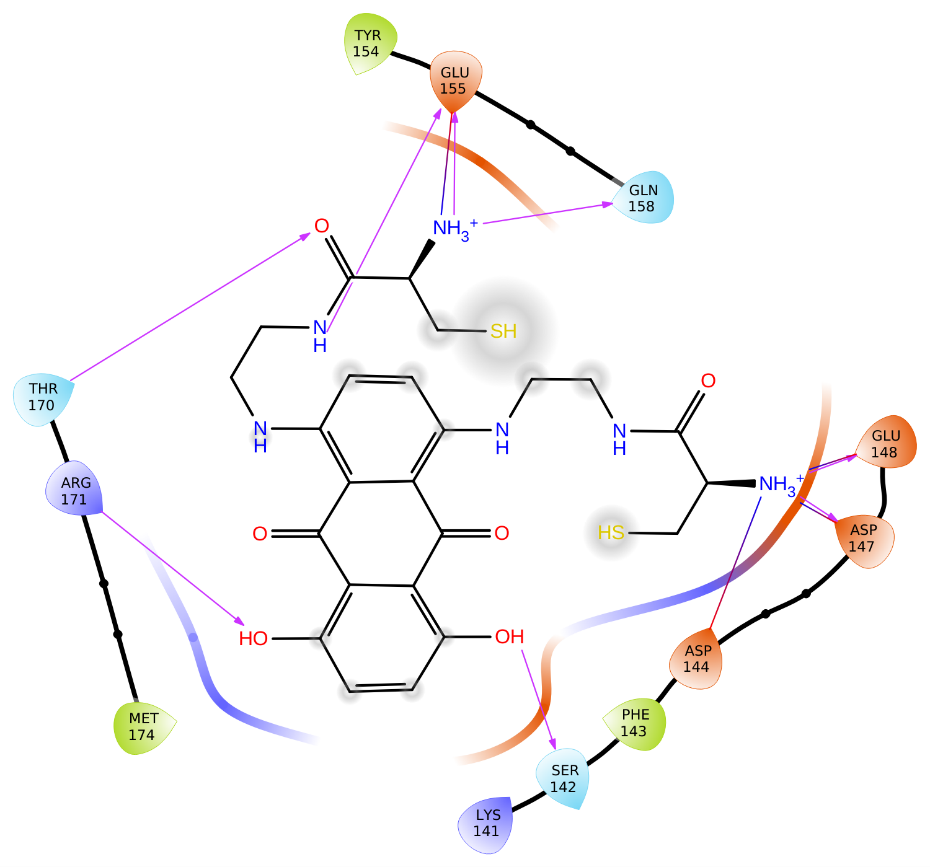

**Supplementary Figure 3:** 2D interaction representation showing interacting residues of NSP1-CTR with highest docking score compound with PubChem ID: 88654295, similar to Mitoxantrone.

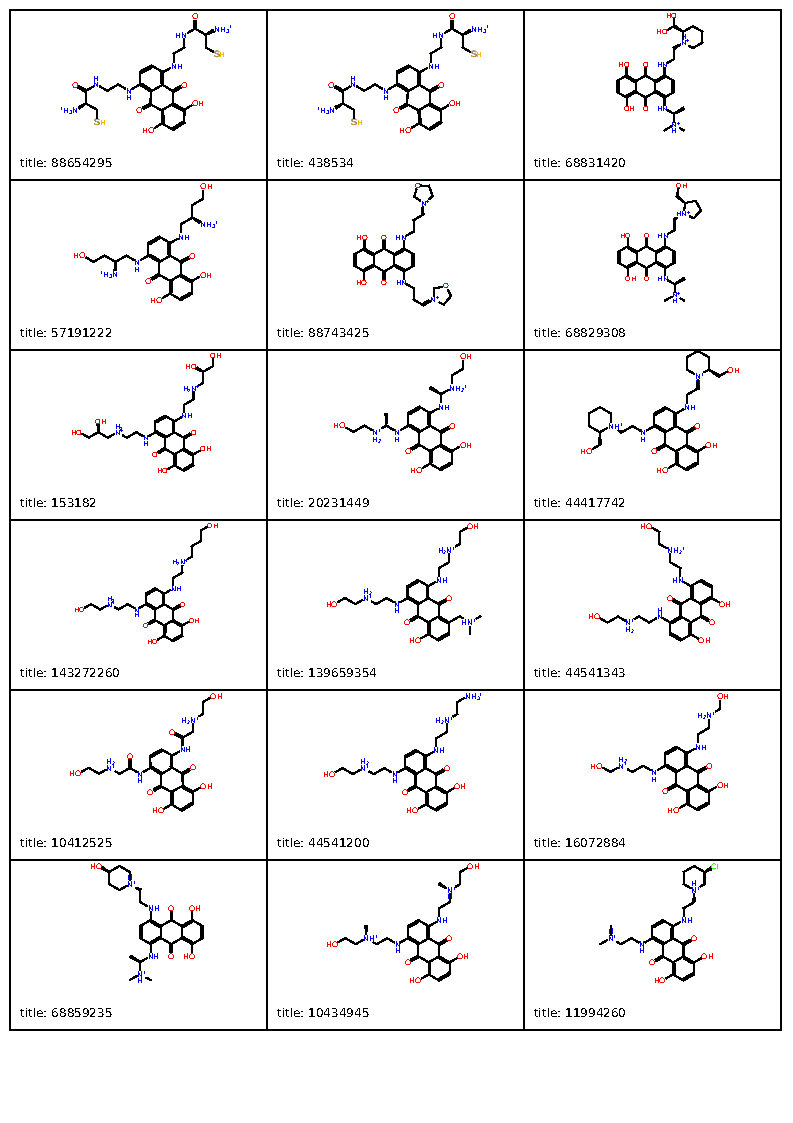

**Supplementary Figure 4A:** Two-dimensional structures of top identified compounds similar to Mitoxantrone from PubChem database with docking score more than -5.3 kcal/mol.

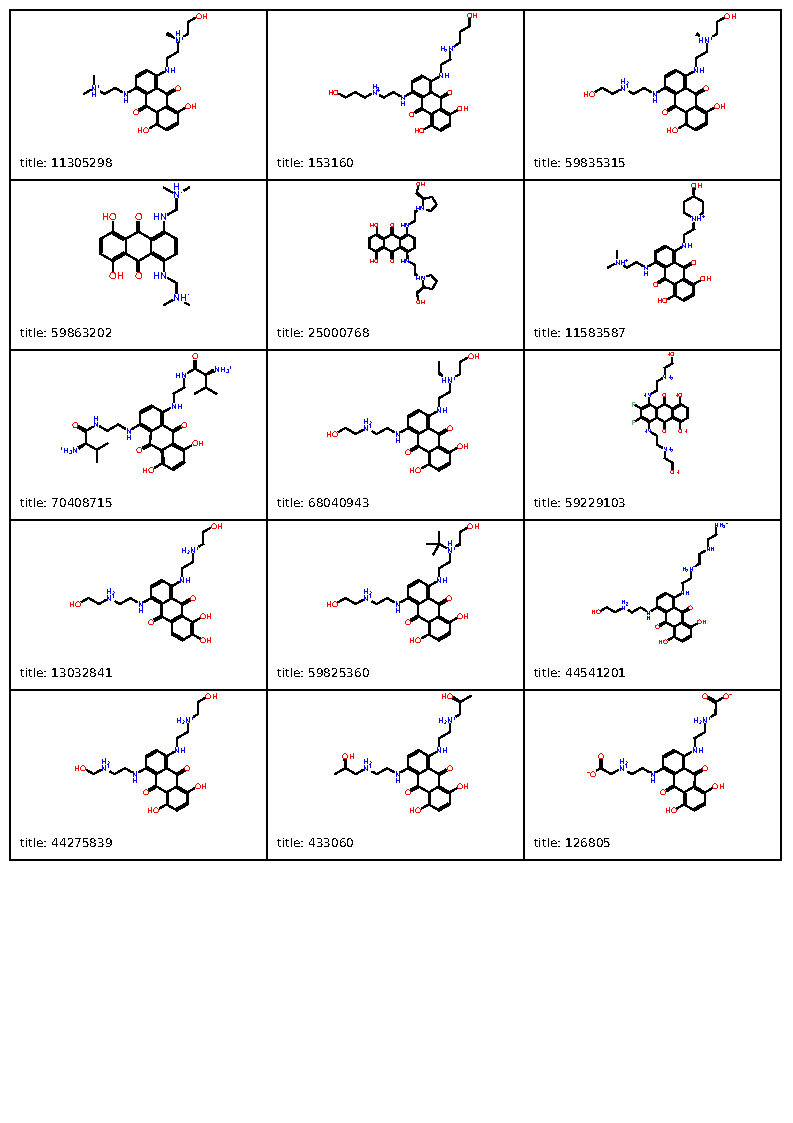

**Supplementary Figure 4B:** Two-dimensional structures of top identified compounds similar to Mitoxantrone from PubChem database with docking score more than -5.3 kcal/mol.

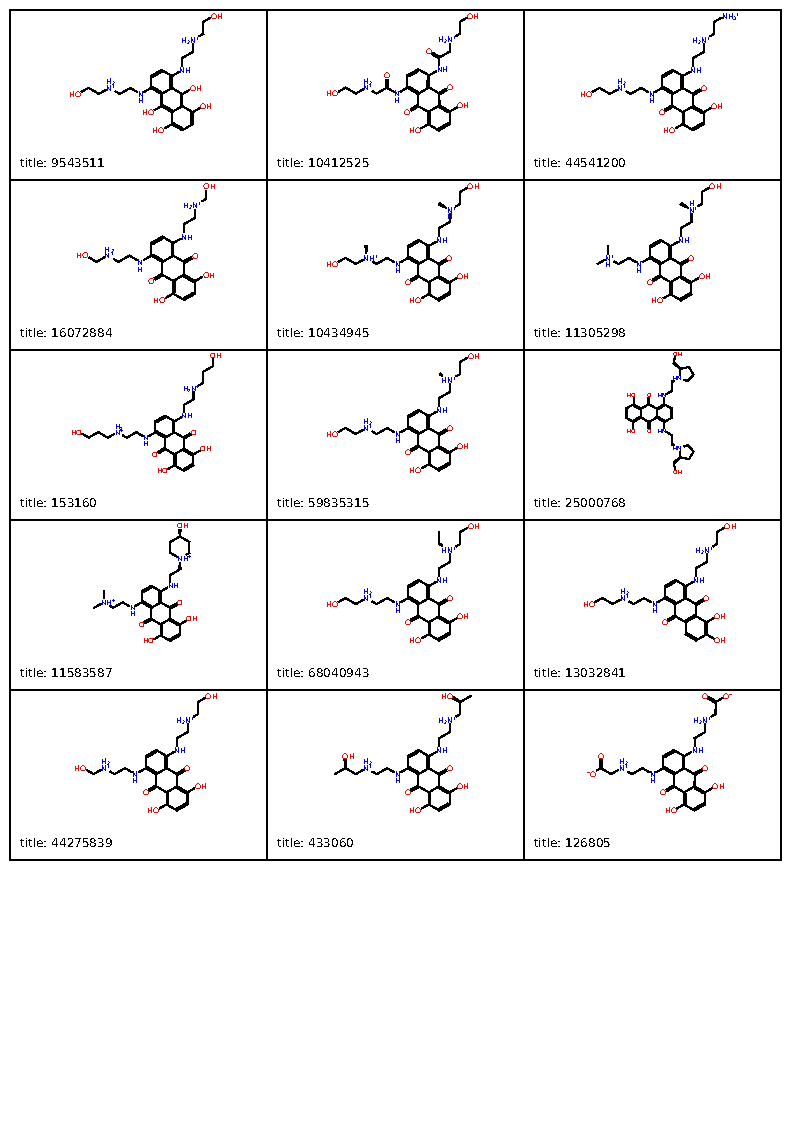

**Supplementary Figure 5:** Two-dimensional structures of top identified conformers similar to Mitoxantrone from PubChem database with docking score more than -5.3 kcal/mol.

**Supplementary Table 1:** Docking scores of top identified compounds similar to Mitoxantrone from PubChem database with docking score more than -5.3 kcal/mol.

| **S. No.** | **PubChem ID** | **Docking score (kcal/mol)** |
| --- | --- | --- |
| 1. | **88654295** | -6.93 |
| 2. | **438534** | -6.93 |
| 3. | **68831420** | -6.66 |
| 4. | **57191222** | -6.34 |
| 5. | **88743425** | -6.17 |
| 6. | **68829308** | -6.10 |
| 7. | **153182** | -6.08 |
| 8. | **20231449** | -6.06 |
| 9. | **44417742** | -6.03 |
| 10. | **143272260** | -6.01 |
| 11. | **139659354** | -5.98 |
| 12. | **44541343** | -5.86 |
| 13. | **10412525** | -5.79 |
| 14. | **44541200** | -5.79 |
| 15. | **16072884** | -5.77 |
| 16. | **68859235** | -5.74 |
| 17. | **10434945** | -5.71 |
| 18. | **11994260** | -5.66 |
| 19. | **11305298** | -5.66 |
| 20. | **153160** | -5.65 |
| 21. | **59835315** | -5.59 |
| 22. | **59863202** | -5.56 |
| 23. | **25000768** | -5.54 |
| 24. | **11583587** | -5.53 |
| 25. | **70408715** | -5.53 |
| 26. | **68040943** | -5.53 |
| 27. | **59229103** | -5.44 |
| 28. | **13032841** | -5.43 |
| 29. | **59825360** | -5.43 |
| 30. | **44541201** | -5.43 |
| 31. | **44275839** | -5.39 |
| 32. | **433060** | -5.38 |
| 33. | **126805** | -5.35 |

**Supplementary Table 2:** Docking scores of top identified conformers similar to Mitoxantrone from PubChem database with docking score more than -5.3 kcal/mol.

| **S. No.** | **PubChem ID** | **Docking score (kcal/mol)** |
| --- | --- | --- |
| 1. | **9543511** | -6.104 |
| 2. | **10412525** | -5.794 |
| 3. | **44541200** | -5.786 |
| 4. | **16072884** | -5.769 |
| 5. | **10434945** | -5.705 |
| 6. | **11305298** | -5.659 |
| 7. | **153160** | -5.647 |
| 8. | **59835315** | -5.589 |
| 9. | **25000768** | -5.535 |
| 10. | **11583587** | -5.534 |
| 11. | **68040943** | -5.527 |
| 12. | **13032841** | -5.426 |
| 13. | **44275839** | -5.389 |
| 14. | **433060** | -5.384 |
| 15. | **126805** | -5.348 |

**Supplementary Movie 1:** Simulation trajectory of one microsecond long NSP1-CTR in complex with MTX using OPLS 2005 forcefield in Desmond simulation package.
